# AttentionAML: An Attention-based Deep Learning Framework for Accurate Molecular Categorization of Acute Myeloid Leukemia

**DOI:** 10.1101/2025.05.20.655179

**Authors:** Lusheng Li, Joseph D. Khoury, Jieqiong Wang, Shibiao Wan

## Abstract

Acute myeloid leukemia (AML) is an aggressive hematopoietic malignancy defined by aberrant clonal expansion of abnormal myeloid progenitor cells. Characterized by morphological, molecular, and genetic alterations, AML encompasses multiple distinct subtypes that would exhibit subtype-specific responses to treatment and prognosis, underscoring the critical need of accurately identifying AML subtypes for effective clinical management and tailored therapeutic approaches. Traditional wet lab approaches such as immunophenotyping, cytogenetic analysis, morphological analysis, or molecular profiling to identify AML subtypes are labor-intensive, costly, and time-consuming. To address these challenges, we propose **AttentionAML**, a novel attention-based deep learning framework for accurately categorizing AML subtypes based on transcriptomic profiling only. Benchmarking tests based on 1,661 AML patients suggested that AttentionAML outperformed state-of-the-art methods across all evaluated metrics (accuracy: 0.96, precision: 0.96, recall of 0.96, F1-score: 0.96, and Matthews correlation coefficient: 0.96). Furthermore, we also demonstrated the superiority of AttentionAML over conventional approaches in terms of AML patient clustering visualization and subtype-specific gene marker characterization. We believe AttentionAML will bring remarkable positive impacts on downstream AML risk stratification and personalized treatment design. To enhance its impact, a user-friendly Python package implementing AttentionAML is publicly available at https://github.com/wan-mlab/AttentionAML.

## Background

Acute myeloid leukemia (AML) is a rapidly progressing hematopoietic malignancy characterized by aberrant clonal expansion of abnormal myeloid progenitor cells, typically leading to bone marrow failure and compromised hematopoiesis (1,2). The heterogeneous nature of AML, manifested through diverse genetic and molecular alterations, presents significant challenges in diagnosis, prognosis, and treatment selection (3,4). Pediatric AML has been categorized into more than 20 molecular subtypes defined by genetic alterations, such as chromosomal alteration, gene fusions, mutations, or tandem duplications (5). These subtypes exhibit varying treatment responses and prognoses, underscoring the critical importance of accurate subtype identification for optimizing therapeutic strategies and improving patient outcomes (6,7). Traditionally, AML subtype identification has relied on a combination of morphological assessment, immunophenotyping, cytogenetics, and molecular profiling (8–10). While these methods have been instrumental in advancing our understanding of AML, the process can be time-consuming, labor-intensive and technically demanding in the clinical practice applications. The technical complexity of these approaches can present significant barriers to their widespread implementation in clinical practice, particularly in resource-limited settings. Moreover, the interpretation of these diverse data types requires significant expertise and can be subject to inter-observer variability.

The advent of the next-generation sequencing (NGS) technologies (11,12), particularly RNA sequencing (RNA-seq), has revolutionized our ability to profile the transcriptome of AML with unprecedented depth and resolution. RNA-seq has emerged as a powerful tool for characterizing the transcriptome of individual tumors, enabling the identification of chromosomal rearrangements and the discovery of novel genetic and clinical markers (13,14). It can provide a comprehensive view of gene expression patterns and reveal critical information about fusion genes that can be critical aspects of AML subtype identification (15–17). However, for fusion-negative subtypes, the classification task becomes more challenging. Unfortunately, mutation calling from RNA-Seq data remains a developing field with inherent limitations for reliable and comprehensive subtype identification (18,19). The challenge is further compounded when dealing with AML subtypes that exhibit similar transcriptional profiles. For instance, subtypes characterized by shared HOX gene expression patterns, such as NPM1, NUP98r, UBTF, DEK::NUP214, and KMT2A-PTD, often present similar transcriptional profiles (5). These similarities of these subtypes pose a significant hurdle for traditional expression-based classification methods, necessitating more sophisticated and comprehensive analytical approaches. In addition, identifying new AML subtypes often involves integrating various NGS techniques, including whole-genome sequencing (WGS) (20), whole-exome sequencing (WES) (21), and RNA-seq. While these approaches provide valuable insights into the genetic and molecular landscape of AML, they are often costly, technically complex, and time-intensive for routine clinical applications. Furthermore, the interpretation of these analyses, such as detecting gene fusions and mutations, requires extensive manual curation and expertise, making the process laborious and susceptible to variability in clinical practice. Recent years, many computational tools based on machine learning algorithms have been developed for the classification of B-cell acute lymphoblastic leukemia (22–26) and T-cell acute lymphoblastic leukemia (27). These advancements have leveraged high-throughput transcriptomic data and machine learning to improve diagnostic accuracy, refine risk stratification models, and inform treatment decisions. In contrast, the application of machine learning to AML is lagging. This gap in knowledge presents a significant opportunity to leverage the power of machine learning to improve the diagnosis, prognosis, and treatment of AML.

To address these challenges, we propose AttentionAML (**Attention**-based Multilayer Perceptron Model for Identifying **A**cute **M**yeloid **L**eukemia Subtypes), a novel approach that leverages the attention mechanisms and multilayer perceptron to accurately classify AML subtypes based solely on transcriptomic profiling. The MLP architecture enhanced with attention mechanisms, which have demonstrated remarkable success in natural language processing tasks, focus on the most relevant features in high-dimensional and complex RNA-seq data. Unlike traditional MLP models, the attention-based MLP captures both global and local dependencies, enhancing the interpretability and performance of the model in subtype classification tasks. This framework allows for effective handling of the high-dimensional nature of RNA-seq data without the need for extensive feature engineering or dimensionality reduction techniques that might lead to loss of important information. Our model is trained on different cohorts of AML patients with known subtypes, utilizing RNA-seq data from publicly available databases such as Therapeutically Applicable Research to Generate Effective Treatments (TARGET) initiative and the St. Jude Cloud platform. In this study, we demonstrated that our AttentionAML Model achieved superior performance in AML subtype identification compared to existing methods in term of accuracy, F1-score and Matthews correlation coefficient. By enabling more accurate and rapid subtype classification, our model aims to address the critical need for precise diagnostic tools in the face of AML’s complex genomic landscape, ultimately contributing to improved patient outcomes and advancing our understanding of this challenging disease.

## Methods

### AML datasets

RNA sequencing data from 1,860 AML patients were analyzed in this study. These datasets comprised patients from multiple cohort genomic studies, including AAML1031 (n=1011) (28), PANAML (n=221) (5), TARGET (n=148) (29), PedAML (n=127) (30), PCGP (n=112) (31,32), RPAML (n=103) (33), SJC-DS-1007 (n=79) (33), TARGET_IF (n=28) (34), G4K (n=24) (35), and CTS (n=7) (36). These RNA-seq datasets were obtained from Target and St. Jude Cloud platform. In the data preprocessing stage, AML patients labeled as “Unclassified” categories were excluded. After this filtering process, our final AML dataset comprised 1,661 patients distributed across 21 distinct molecular subtypes (Fig. 1A). A detailed breakdown of the dataset composition by subtype is illustrated in the pie chart (Fig. 1A), highlighting significant imbalances among categories. For instance, subtypes such as KMT2Ar (26%), RUNX1::RUNX1T1 (16%) and CBFB::MYH11 (13%) accounted for a large proportion of the dataset compared to rarer subtypes like MNX1 and RUNX1::RUNX1T1-like. The age distribution of patients in the AML dataset is illustrated in Fig. 1B. The histogram reveals two distinct age groups, categorized as “Childhood” (patients aged 0– 15 years, shown in red) and “Adolescent and Young Adults (AYA)” (patients aged 16–40 years, shown in blue). The majority of cases are concentrated within the Childhood group, with a prominent peak in patients under the age of 5. As age increases, the frequency of cases gradually decreases, with fewer cases observed in the AYA group. This distribution highlights a higher incidence of the condition in younger patients, aligning with the expected age-related epidemiological characteristics.

**Figure 1.**
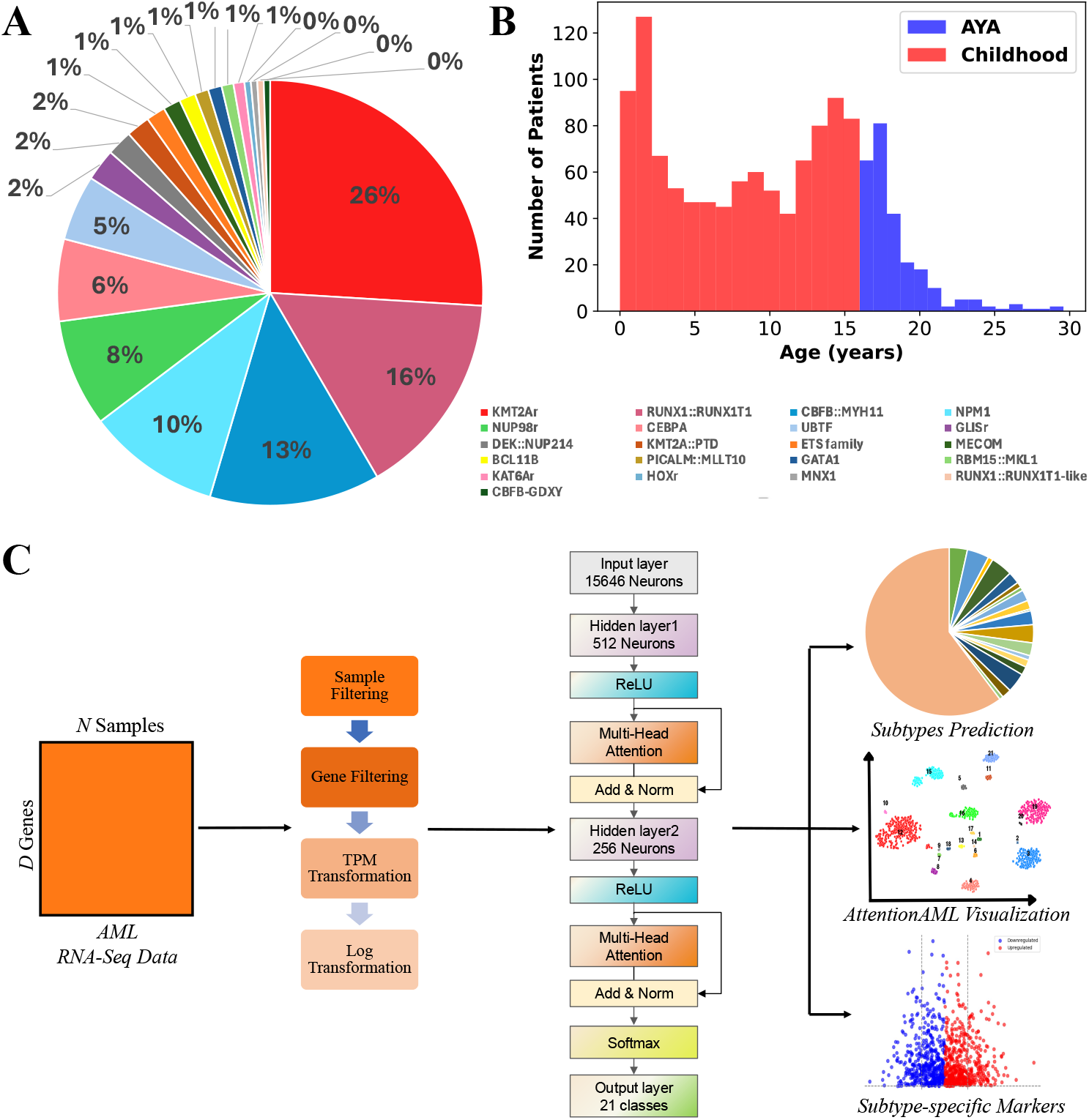
Overview of AML subtype identification study using AttentionAML framework. **(A)** The breakdown of the AML dataset. The pie chart showing the subtype distribution of 1,860 AML patients across 21 distinct categories. The percentages indicate the relative prevalence of each subtype within the AML dataset. **(B)** The age distribution of the AML dataset. Histogram illustrating the age distribution of AML patients, categorized into childhood (red) and adolescent and young adult (AYA, blue) groups. **(C)** Workflow of the AttentionAML model for AML subtype classification. The workflow begins with preprocessing steps applied to AML RNA-seq data, including sample filtering, gene filtering, transcripts per million (TPM) normalization, and log transformation. Subsequently, processed data serve as input to the AttentionAML neural network model. The final outputs include subtype prediction probabilities, visualization of subtypes, and identification of subtype-specific gene markers.

### AttentionAML framework

We implemented a multi-head attention-based MLP for AML subtype classification. The architecture of our proposed AttentionAML model (Fig. 1C) is composed of several key layers designed to efficiently capture both global and local dependencies within high-dimensional gene expression data. The architecture consists of an input layer with 15,646 neurons, corresponding to the gene features in the dataset, followed by two hidden layers with multi-head attention mechanisms (37). The first hidden layer contains 512 neurons, and the second has 256 neurons. Each hidden layer is followed by a ReLU activation function to introduce non-linearity into the model (38). Multi-head attention modules are incorporated after each hidden layer to capture complex interactions between features. These self-attention mechanisms use query, key, and value transformations, each implemented as linear layers with 512 and 256 features for the first and second attention modules, respectively. The multi-head attention block outputs to an add & norm layer, where the residual connections are combined with the input and then normalized, promoting stable and efficient training. The final layers of the model consist of a softmax layer, which outputs a probability distribution over 21 classes, corresponding to the number of AML subtypes. With the gene expression data of AML patients as input, the proposed AttentionAML framework demonstrates compatibility by accepting multiple formats of gene expression data, including raw counts, Fragments Per Kilobase of transcript per Million mapped reads (FPKM), and Transcripts Per Million (TPM). All expression data are standardized through transformation to log2(TPM + 1) values to ensure consistency for AML subtype prediction. The AttentionAML framework consisted of three main steps: (1) data preprocessing and normalization, (2) multi-head attention-based MLP, and (3) multi-class classification, as outlined in Fig. 1C. This framework allows for effective handling of the high-dimensional RNA-seq data without the need for extensive feature engineering or dimensionality reduction techniques that might lead to loss of important information. It enables precise classification of AML subtypes, improved visualization and identification of subtype-specific biomarkers.

### Data preprocessing

RNA-seq data from 1,860 AML patients were preprocessed illustrated as Fig. 1C to ensure data quality and comparability. Initially, raw read counts were obtained from different cohort datasets. The patients with “unclassified” subtype were excluded, resulting in 1,661 patients being retained for further preprocessing. For the gene filtering, only the gene expressed in at least 75% of the samples in each dataset were retained and the common gene list from each dataset were selected, yielding a set of 15,646 genes. This filtering step removed rarely expressed genes that could potentially introduce noise and reduce the robustness of downstream analyses. Subsequently, raw read counts were converted to Transcripts Per Million (TPM). This step accounts for both sequencing depth and gene length, enabling biologically meaningful comparisons across samples (39). To address the inherent skewness of gene expression data, TPM values were transformed using the formula log2(TPM+1). The resulting log-transformed TPM values were utilized for subsequent model training and AML subtype identification.

### Subtype-specific differential gene expression analysis

For subtype-specific differential gene expression (DGE) analysis, we utilized the expected raw counts from AML samples. Low-abundance genes were removed using a CPM threshold corresponding to 10 read counts. The trimmed mean of M-values (TMM) method (40) was employed to calculate normalization factors, followed by variance stabilization using the voom transformation. Differential expression analysis was performed on the normalized counts using the lmFit and eBayes functions from the limma R package v3.54.2 (41). Significantly differentially expressed genes were identified using stringent criteria of FDR < 0.05 and |log2FC| > 1. Heatmap of expression patterns across subtypes was generated using the Pheatmap package (1.0.12) (42).

### Performance Evaluation

To rigorously assess the performance of our AttentionAML model for AML subtype classification, we employed a comprehensive set of evaluation metrics, including Accuracy, Precision, Recall, F1-Score (F1), and the Matthews Correlation Coefficient (MCC). These complementary metrics are defined as:

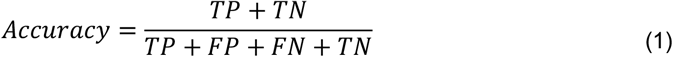

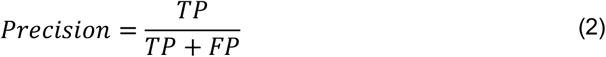

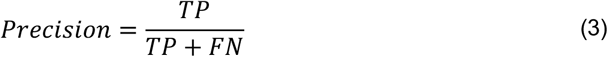

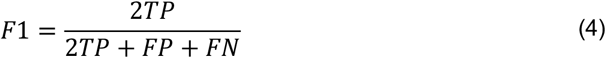

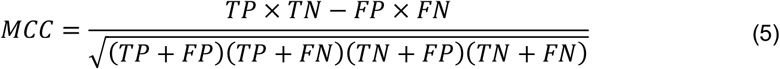

where TP, TN, FP, and FN represent true positives, true negatives, false positives, and false negatives, respectively. Accuracy is calculated as the ratio of correctly predicted samples to the total number of samples, providing a general measure of the model’s ability to classify AML subtypes correctly. Precision, defined as the number of true positives divided by the sum of true positives and false positives, measures the model’s ability to avoid false positives, which is critical when distinguishing between similar AML subtypes. Recall, calculated as the ratio of true positives to the sum of true positives and false negatives, evaluates the model’s capacity to identify all positive samples. The F1-Score represents the harmonic mean of precision and recall, providing a balanced metric that accounts for both false positives and false negatives, especially useful in imbalanced datasets. Additionally, we computed the MCC, a metric that takes into account true positives, true negatives, false positives, and false negatives, offering a comprehensive measure of classification quality. It is particularly useful in multi-class classification tasks with imbalanced classes. These metrics were calculated using scikit-learn’s metrics module, with macro-averaging applied for multi-class scenarios to treat all classes equally. Performance was evaluated using 100 times stratified k-fold cross-validation (k=5) to ensure robust estimates across all AML subtypes.

## Results

### Enhancing Model Performance through Attention Mechanisms

To evaluate the performance of the AttentionAML model and its potential generalizability to new data, we utilized a robust 5-fold cross-validation repeated 100 times. This was conducted on an RNA-seq dataset containing 1,661 AML samples across 21 subtypes, as outlined in Fig. 1A. The figure illustrated the performance comparison of three models: Ensemble SVM, MLP, and AttentionAML, for AML subtype classification. Ensemble SVM was adopted from RanBALL, an ensemble Random projection-based model for identifying B-ALL subtypes (26). Across all five key performance metrics, AttentionAML consistently outperformed both Ensemble SVM and MLP models. The AttentionAML model produced impressive average outcomes, achieving an accuracy of 96.53%, a precision of 96.54%, a recall of 96.53%, an F1 score of 96.38%, and a MCC of 0.96 (Fig. 2A). The superior F1-score achieved by AttentionAML indicated a more optimal balance between precision and recall compared to existing state-of-the-art models. Furthermore, the MCC provided a robust performance metric, particularly suited for scenarios involving imbalanced class distributions. The MCC results highlighted AttentionAML’s ability to maintain high classification accuracy across all subtypes, including those with limited sample sizes. Notably, it achieved superior median scores and exhibited less variability, as evidenced by the compact interquartile ranges in the box plots. These results indicates that the attention mechanism incorporated into the AttentionAML model improves its ability to distinguish between AML subtypes more effectively.

**Figure 2.**
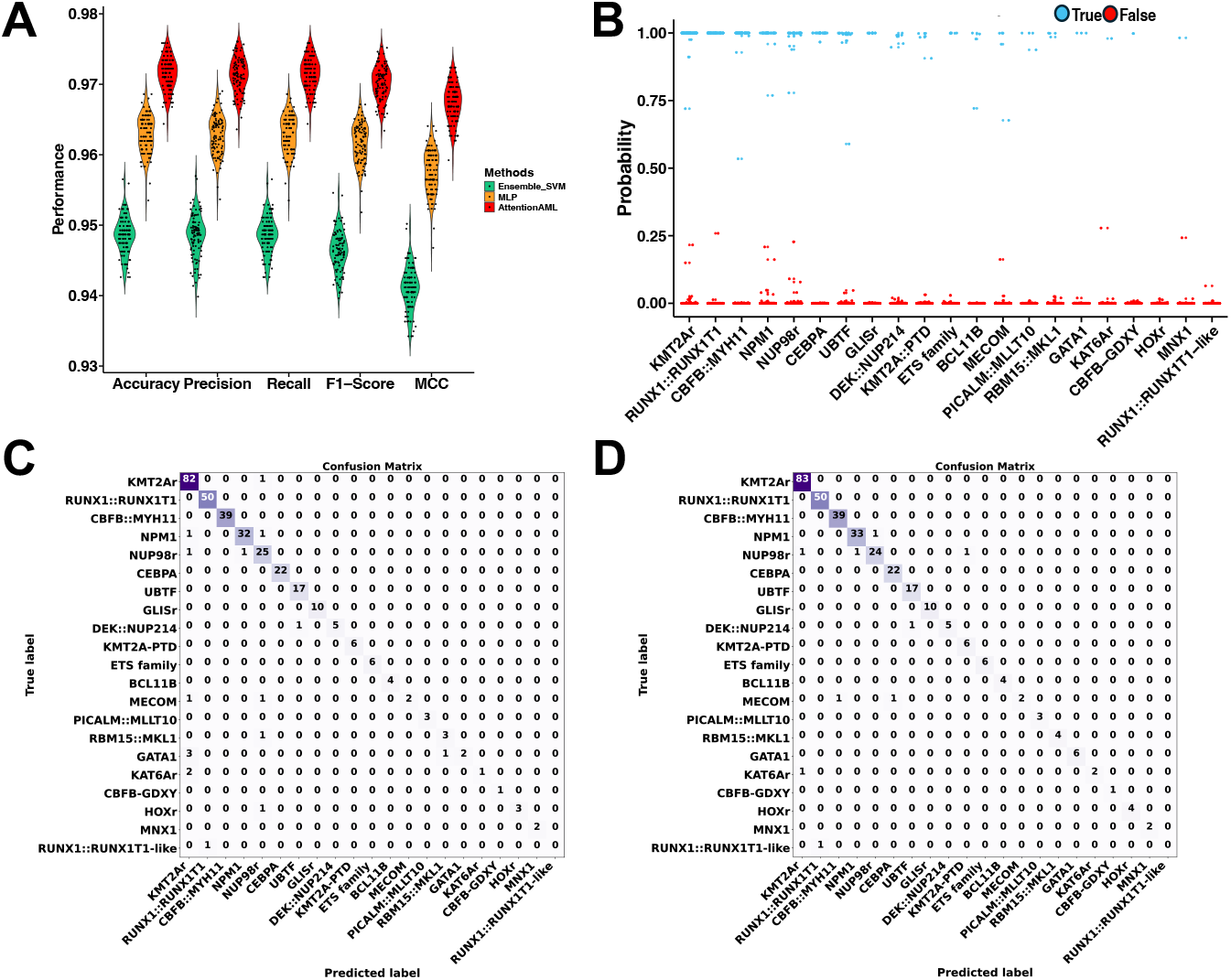
Performance comparison of AttentionALL and state-of-the-art methods for AML subtyping. **(A)** The performance of Ensemble_SVM, MLP, and AttentionAML on multiple evaluation metrics across 100 iterations of 5-fold cross-validation. **(B)** Prediction probability distribution for the 20% held-out test set using AttentionAML. Each point represents the probability of a sample (out of 332) being classified into a specific AML subtype. The blue dots indicate the subtype predicted by AttentionAML model, aligned with the categories on the horizontal axis. **(C, D)** Confusion matrices comparing the 20% held-out test set performance of the AttentionAML **(C)** and Ensemble_SVM **(D)** models. Each matrix element shows the number of samples classified, with the diagonal representing correct classifications (True Positives). Color intensity correlates with the number of samples.

To further validate our AttentionAML model, we evaluated its performance on an independent hold-out test set derived from the AML dataset. This test set comprised 332 patients, representing 20% of the total AML datset, randomly selected to ensure unbiased assessment. The AttentionAML model exhibited robust performance, achieving an accuracy of 97.92% on this held-out subset. Fig. 2B illustrated the prediction probabilities for each test sample, revealing the model’s consistent ability to generate high-confidence predictions across various AML subtypes. The probabilities distribution predominantly clustered near a probability of 1.0, underscores the model’s proficiency in capturing intricate gene expression patterns specific to each AML subtype. This high-confidence output not only demonstrates the model’s accuracy but also suggests its potential reliability in clinical applications, where confident predictions are crucial for informed decision-making. The consistent high-confidence predictions across diverse AML subtypes highlight the AttentionAML model’s potential as a valuable tool for precise and reliable AML subtype classification in clinical settings. The 20% held-out test was also performed with the Ensemble SVM and MLP methods. The AttentionAML model demonstrates superior performance across all metrics, achieving an accuracy of 97.92% compared to 94.12% for the Ensemble SVM. Notably, our model shows marked improvements in precision (97.28% vs. 93.53%), recall (97.92% vs. 94.12%), F1 score (97.50% vs. 93.55%), and Matthews Correlation Coefficient (97.60% vs. 93.20%). These results underscore the robust and balanced performance of the AttentionAML model across various AML subtypes. The confusion matrices provide further insight into the classification performance of both models (Fig. 2C-D). The AttentionAML model exhibits more diagonal elements, indicating higher correct classifications across most subtypes. Particularly noteworthy is the enhanced performance in identifying rarer subtypes such as BCL11B, MECOM, and PICALM::MLLT10, where the Ensemble SVM showed more misclassifications. The AttentionAML model also demonstrates reduced off-diagonal elements, signifying fewer misclassifications between different AML subtypes. These results highlight the AttentionAML model’s enhanced ability to capture complex gene expression patterns specific to each AML subtype, leading to more accurate and reliable classifications. This improved performance across a diverse range of AML subtypes suggests the potential of our model as a valuable tool for precise AML diagnosis and subtype classification in clinical settings.

### AttentionAML outperforms t-SNE for high-dimensional data visualization

Leveraging embedding information derived from the hidden layer of AttentionAML model, AttentionAML outperformed state-of-the-art visualization techniques. Specifically, the 1024-dimension embedding was extracted and projected into two-dimensional space for visualization using t-SNE. It demonstrated superior capability in revealing subtype-specific patterns and maintaining the integrity of the underlying data structure, providing a more accurate and interpretable representation of the complex relationships between AML subtypes. To benchmark its performance, we utilized t-SNE, a widely recognized and powerful method for visualizing high-dimensional data.

Fig. 3A illustrated the AttentionAML’s capability to visualize the AML samples, where distinct subtypes form well-defined clusters, underscoring the model’s ability to preserve and emphasize the inherent structure of the data. This visualization facilitates the identification and interpretation of 21 unique subtypes, spanning from prevalent subtypes like KMT2Ar and RUNX1::RUNX1T1 to less common ones such as the MNX1 and RUNX1::RUNX1T1-like. Each data point represents an individual sample, colored and labeled according to its corresponding molecular subtype. Distinct clusters indicate that the model successfully captures subtype-specific features, demonstrating its capacity to separate subtypes based on underlying molecular patterns. Notably, subtypes with genetic similarities, such as CBFB-GDXY and CBFB::MYH11, along with RUNX1::RUNX1T1 and its RUNX1::RUNX1T1-like variant, cluster in close proximity to each other, indicating their shared molecular characteristics in the latent space. The clear separation between different AML subtypes underscored the robustness of the model in learning discriminative features. Importantly, the embedding space highlights clusters of rarer subtypes, suggesting the model’s sensitivity to subtle differences even among underrepresented classes. In contrast, Fig. 3B displayed a t-SNE visualization that lacks the embedding information. It exhibited reduced structural clarity and increased scatter in the data distribution, with less defined boundaries between AML subtypes. AML subtypes such as CBFB-GDXY, CBFB::MYH11NPM1, KMT2Ar, and NUP98r showed substantial overlap and poor cluster boundary. The significant contrast between these visualizations highlights the effectiveness of our model’s approach in generating meaningful and interpretable representations of complex transcriptomic profiles in AML. By leveraging embedding information, this visualization technique not only effectively captures unique molecular patterns associated with each subtype but also potentially reveals biologically meaningful relationships between subtypes, ultimately advancing our understanding of disease heterogeneity or molecular mechanisms.

**Figure 3.**
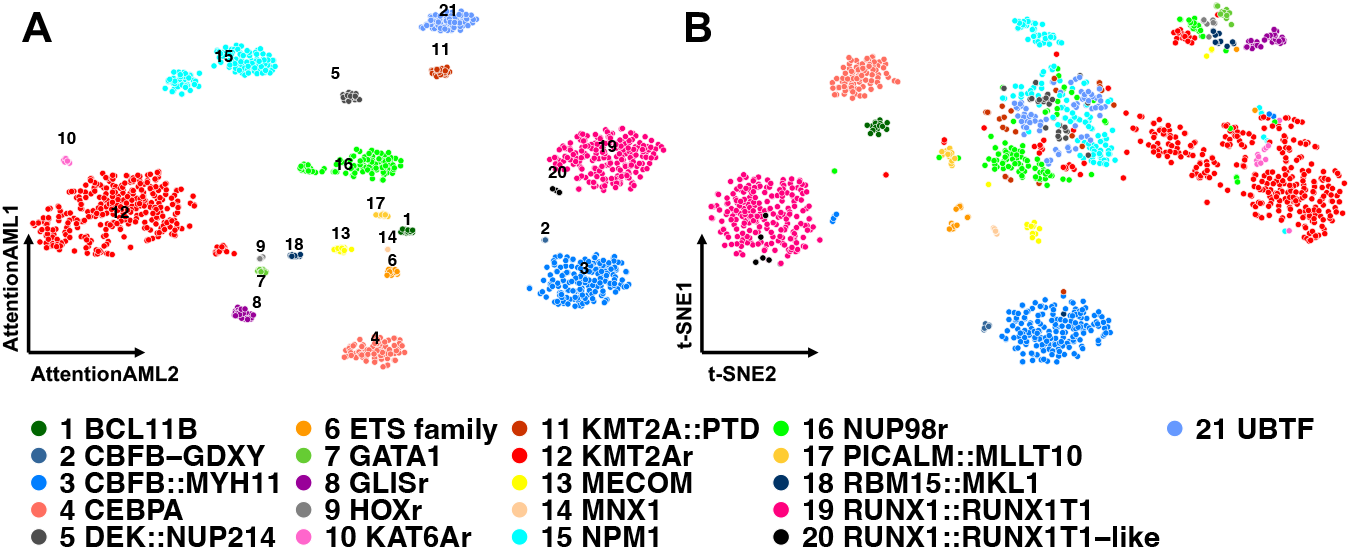
Comparing AttentionAML and t-SNE visualizations of AML subtypes. **(A)** AttentionAML visualization of MLP hidden layer embedding information. **(B)** t-SNE visualization of the reduced dimension matrix with conventional gene expression profiling information only. Each point represents a sample, with distinct colors denoting different AML subtypes. The same color scheme was used in both plots.

### Subtype-specific differential gene expression analysis of AML patients

The subtype-specific differential gene expression analysis for each AML subtype was conducted to identify and explore the subtype-specific biomarkers. The heatmap visualization (Fig. 4A) clearly illustrated distinct subtype-specific gene expression patterns, highlighting key marker genes that are significantly upregulated in particular AML subtypes relative to others. Volcano plots (Fig. 4B–U) showed significantly upregulated (red-labeled) and downregulated (blue-labeled) genes unique to each AML subtype, clearly defining molecular heterogeneity among these subgroups. The KMT2Ar subtype (Fig. 4B) displayed enrichment of genes including LAMP5, HOXA9, HOXA3, ZNF521 and MEIS1. LAMP5 is upregulated in certain high-risk fusion subtypes of AML, including KMT2Ar, KAT6Ar, and NUP98r (43). Similarly, the CEBPA subtype (Fig. 4G) also showed upregulation of gene LAMP5. HOXA9 is often overexpressed in AML and is associated with poor prognosis (44). HOXA3 and MEIS1 are key regulator in hematopoiesis (45,46). ZNF521 is a transcription co-factor with regulatory functions in haematopoietic cells (47). The RUNX1::RUNX1T1 subtype (Fig. 4C) exhibited significant upregulation of transcription factors implicated in hematopoiesis regulation. In the CBFB::MYH11 and RBM15::MKL1 subtypes (Fig. 4D,P), notable upregulation of genes such as MYH11 and MN1 were observed. MN1 can be rearranged and overexpressed, leading to a poor prognosis (48). The BCL11B subtype (Fig. 4M) exhibited significant upregulation of BCL11B itself and other transcription factors regulated in hematopoiesis. The NPM1, NUP98r, UBTF, DEK::NUP214, PICALM::MLLT10 and KAT6Ar subtypes (Fig. 4E, F, H, J, O, R) also showed significant overexpression of HOXA cluster genes, consistent with prior knowledge linking these genes to AML prognosis. Moreover, other subtypes such as GLISr (Fig. 4I), ETS family (Fig. 4L), MECOM (Fig. 4N), GATA1 (Fig. 4Q) and MNX1 (Fig. 4U) demonstrated strong differential expression of genes associated with myeloid differentiation and cellular proliferation, reflecting its known pathogenic mechanisms.Subtypes characterized by specific genetic rearrangements, such as KMT2A::PTD (Fig. 4K), CBFB-GDXY (Fig. 4S) and HOXr (Fig. 4T), similarly showcased distinct gene signatures emphasizing their unique oncogenic drivers. These analyses provide critical insights into the molecular characteristics defining AML subtypes, offering potential biomarkers for refined diagnostic stratification and targeted therapeutic strategies tailored to specific AML molecular profiles.

**Figure 4.**
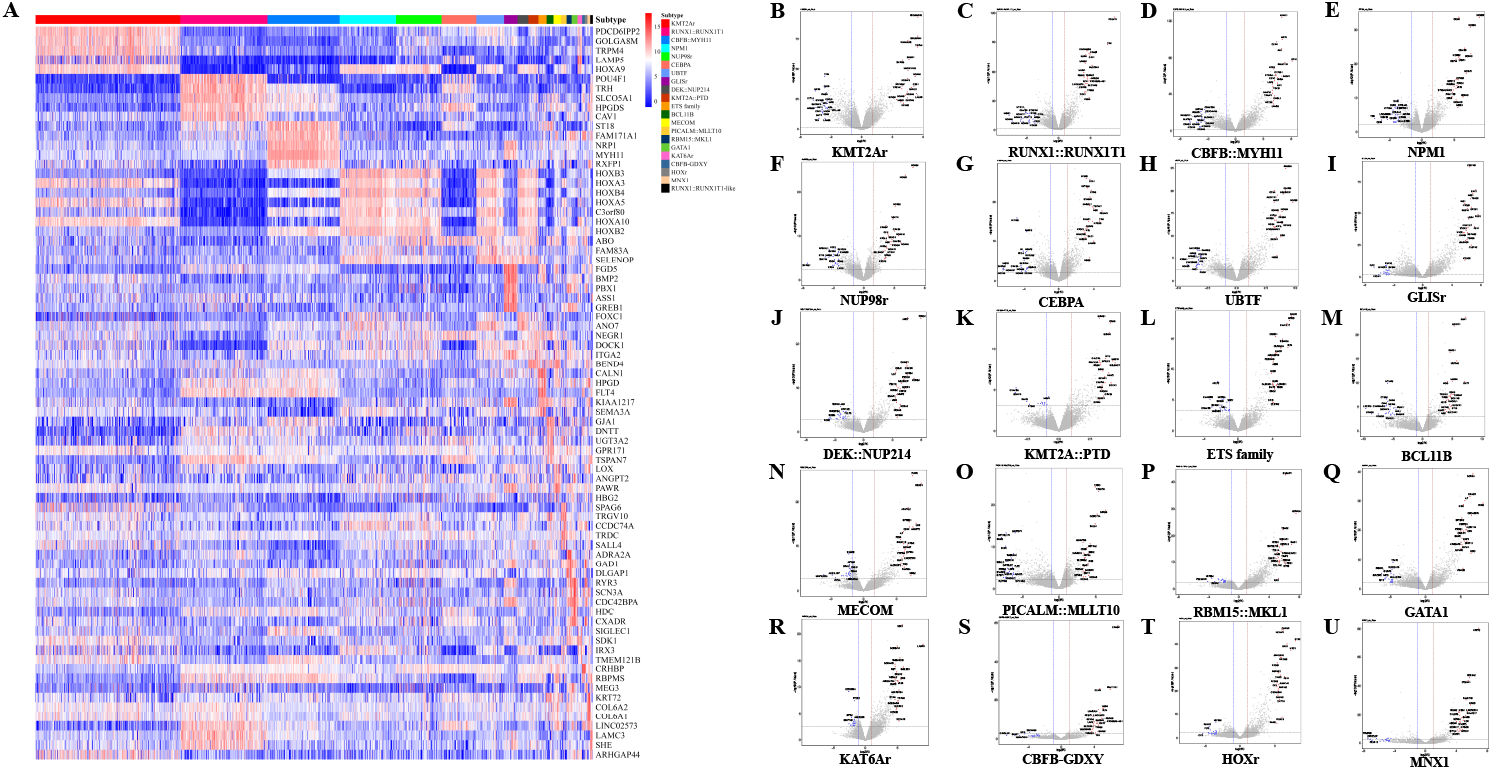
Subtype-specific differential gene expression analysis within AML subtypes. **(A)** Heatmaps displayed expression patterns of the top 5 DEGs for each subtype comparison. **(B∼U)** The volcano plot of DEGs for each of the 20 AML subtypes. Each dot represents a gene, with significantly upregulated genes (log2 fold change ≥ 1, adjusted p-value < 0.05) marked in red, and significantly downregulated genes (log2 fold change ≤ -1, adjusted p-value < 0.05) marked in blue. Notable genes demonstrating strong subtype-specific expression are labeled explicitly. Horizontal dashed line represents the significance threshold (adjusted p-value < 0.05), while vertical dashed lines denote thresholds for fold change (log2 fold change = ±1).

## Discussion

In this study, we presented AttentionAML, an innovative attention-based MLP model, for accurately classifying acute myeloid leukemia (AML) subtypes using transcriptomic profiling only. By leveraging attention mechanisms, the model effectively captured complex interactions between gene features, allowing for a more precise classification of AML subtypes. In addition, the multi-head attention mechanism proves crucial in capturing intricate relationships within high-dimensional transcriptomic data. Unlike traditional machine learning approaches, our model can effectively process RNA-seq data without extensive feature engineering, preserving critical molecular information that might be lost through conventional dimensionality reduction techniques. AttentionAML achieved superior performance compared to state-of-the-art methods, such as Ensemble SVM and MLP approaches, as demonstrated by consistently high metrics across accuracy, precision, recall, F1-score, and MCC. Notably, it exhibited superior performance in correctly classifying rare and underrepresented subtypes, such as BCL11B and MECOM, which have traditionally posed significant challenges for classification models.

In addition to accurate subtype classification, AttentionAML demonstrated superior visualization capabilities, outperforming traditional t-SNE-based methods by leveraging embedding information from the MLP hidden layers. Compared to traditional t-SNE techniques, AttentionAML’s embedding visualization revealed more distinct and interpretable clusters of AML subtypes. This enhanced visualization method offers insights into the molecular landscape of AML, potentially uncovering previously unrecognized relationships between subtypes. The visualization highlighted nuanced distinctions between subtypes, even those with genetic similarities, such as, subtypes like CBFB-GDXY and CBFB::MYH11, or RUNX1::RUNX1T1 and its RUNX1::RUNX1T1-like variant. This level of identificaion is critical for precise diagnostic and therapeutic strategies.

The AttentionAML framework offers transformative potential for clinical oncology by addressing critical challenges in AML diagnosis and management. By providing a rapid, accurate, and reproducible method for subtype classification, the model can significantly reduce diagnostic complexity and inter-observer variability inherent in traditional morphological and cytogenetic approaches. The high-confidence predictions, with probabilities consistently near 1.0, suggest strong potential for clinical translation, enabling more precise risk stratification and personalized treatment strategies. Particularly impactful is the model’s ability to effectively classify rare AML subtypes, which often pose significant diagnostic challenges. Moreover, by capturing molecular biomarkers between subtypes, AttentionAML could accelerate our understanding of AML’s complex genomic landscape, ultimately supporting the development of targeted therapeutic interventions and improving patient outcomes across diverse pediatric AML populations.

While AttentionAML demonstrated significant promise, several limitations and future research opportunities exist. The current model requires extensive validation across larger and more diverse patient cohorts to ensure generalizability. The publicly available datasets may introduce potential biases, such as overrepresentation of specific subtypes or technical variability between datasets (49). The batch effect from different datasets could be mitigated by computational methods, such as ComBat (50). In addition, the inherent imbalance among AML subtypes within the dataset presents a potential challenge for model performance and generalizability. Minority subtypes can potentially introduce bias and reduce the model’s ability to accurately classify less frequent molecular subtypes. By synthetically expanding the representation of minority subtypes, we can enhance the model’s learning capabilities and improve its robustness across diverse molecular subtypes. Techniques such as generative adversarial networks (GANs) (51), synthetic minority oversampling technique (SMOTE) (52), or advanced deep learning-based augmentation methods (53) can generate synthetic molecular representations. Furthermore, integrating multi-omics data, such as genetics, epigenomics or imaging (54), with transcriptomic information could enhance the model’s ability to capture complex biological interactions and further improve its predictive accuracy.

The development of our computational framework holds transformative potential for AML clinical management, offering direct positive impacts on diagnostic precision, personalized treatment strategies, and risk stratification. The critical significance lies in the differential treatment responses and survival rates associated with distinct AML subtypes. By enabling precise molecular subtype identification, our approach empowers clinicians to select the most targeted and effective treatment strategies tailored to individual patient profiles. To facilitate further research, a Python package implementing AttentionAML is publicly available on GitHub (https://github.com/wan-mlab/AttentionAML).

## Acknowledgements

The authors would like to express our gratitude to St. Jude Cloud platform (https://www.stjude.cloud) and TARGET, which provided publicly accessible genomic data. Special thanks to all members of Dr. Wan’s lab for insightful discussions.

## Authors’ contributions

L.L.: data preprocessing, machine learning model development, data analysis and interpretation, manuscript preparation, editing, and review. J.D.K.: manuscript editing and review. J.W.: manuscript editing and review. S.W.: study concept and design, manuscript editing and review.

## Data availability

The RNA-seq data of AML samples can be publicly accessed from St. Jude Cloud and TARGET. The AttentionAML package can be accessed at https://github.com/wan-mlab/AttentionAML.

## Reference

1. Klco JM, Spencer DH, Miller CA, Griffith M, Lamprecht TL, O’Laughlin M, et al. Functional heterogeneity of genetically defined subclones in acute myeloid leukemia. Cancer Cell. 2014;25(3):379–92.

2. Khwaja A, Bjorkholm M, Gale RE, Levine RL, Jordan CT, Ehninger G, et al. Acute myeloid leukaemia. Nat Rev Dis Primer. 2016;2(1):1–22.

3. Lagunas-Rangel FA, Chávez-Valencia V, Gómez-Guijosa MÁ, Cortes C. Acute Myeloid Leukemia—Genetic Alterations and Their Clinical Prognosis.. Volume. 11(4).

4. De Kouchkovsky I, Abdul-Hay M. ‘Acute myeloid leukemia: a comprehensive review and 2016 update.’ Blood Cancer J. 2016 Jul 1;6(7):e441–e441.

5. Umeda M, Ma J, Westover T, Ni Y, Song G, Maciaszek JL, et al. A new genomic framework to categorize pediatric acute myeloid leukemia. Nat Genet. 2024 Feb;56(2):281–93.

6. Marcucci G, Haferlach T, Döhner H. Molecular Genetics of Adult Acute Myeloid Leukemia: Prognostic and Therapeutic Implications. J Clin Oncol. 2011 Feb 10;29(5):475–86.

7. Watts J, Nimer S. Recent advances in the understanding and treatment of acute myeloid leukemia. F1000Research. 2018 Aug 6;7:1196.

8. Moarii M, Papaemmanuil E. Classification and risk assessment in AML: integrating cytogenetics and molecular profiling. Hematol 2014 Am Soc Hematol Educ Program Book. 2017;2017(1):37–44.

9. Marceau-Renaut A, Duployez N, Ducourneau B, Labopin M, Petit A, Rousseau A, et al. Molecular profiling defines distinct prognostic subgroups in childhood AML: a report from the French ELAM02 study group. HemaSphere. 2018;2(1):e31.

10. Bain BJ, Béné MC. Morphological and Immunophenotypic Clues to the WHO Categories of Acute Myeloid Leukaemia. Acta Haematol. 2019 Apr 9;141(4):232–44.

11. Behjati S, Tarpey PS. What is next generation sequencing? Arch Dis Child-Educ Pract. 2013;

12. Dahui Qin. Next-generation sequencing and its clinical application. Cancer Biol Med. 2019 Feb 1;16(1):4.

13. Byron SA, Van Keuren-Jensen KR, Engelthaler DM, Carpten JD, Craig DW. Translating RNA sequencing into clinical diagnostics: opportunities and challenges. Nat Rev Genet. 2016 May 1;17(5):257–71.

14. Tsimberidou AM, Fountzilas E, Bleris L, Kurzrock R. Transcriptomics and solid tumors: The next frontier in precision cancer medicine. Precis Med Cancer. 2022 Sep 1;84:50–9.

15. Valk Peter J.M., Verhaak Roel G.W., Beijen M. tAntoinette, Erpelinck Claudia A.J., van Doorn-Khosrovani Sahar Barjesteh van Waalwijk, Boer Judith M., et al. Prognostically Useful Gene-Expression Profiles in Acute Myeloid Leukemia. N Engl J Med. 350(16):1617– 28.

16. Arindrarto W, Borràs DM, de Groen RAL, van den Berg RR, Locher IJ, van Diessen SAME, et al. Comprehensive diagnostics of acute myeloid leukemia by whole transcriptome RNA sequencing. Leukemia. 2021 Jan 1;35(1):47–61.

17. Kerbs P, Vosberg S, Krebs S, Graf A, Blum H, Swoboda A, et al. Fusion gene detection by RNA-sequencing complements diagnostics of acute myeloid leukemia and identifies recurring NRIP1-MIR99AHG rearrangements. Haematologica. 2022 Jan 1;107(1):100–11.

18. Kaya C, Dorsaint P, Mercurio S, Campbell AM, Eng KW, Nikiforova MN, et al. Limitations of Detecting Genetic Variants from the RNA Sequencing Data in Tissue and Fine-Needle Aspiration Samples. Thyroid®. 2021 Apr 1;31(4):589–95.

19. Abbasi A, Alexandrov LB. Significance and limitations of the use of next-generation sequencing technologies for detecting mutational signatures. DNA Repair. 2021 Nov 1;107:103200.

20. Ng PC, Kirkness EF. Whole Genome Sequencing. In: Barnes MR, Breen G, editors. Genetic Variation: Methods and Protocols [Internet]. Totowa, NJ: Humana Press; 2010. p. 215–26. Available from: 10.1007/978-1-60327-367-1_12

21. Rabbani B, Tekin M, Mahdieh N. The promise of whole-exome sequencing in medical genetics. J Hum Genet. 2014;59(1):5–15.

22. Mäkinen VP, Rehn J, Breen J, Yeung D, White DL. Multi-Cohort Transcriptomic Subtyping of B-Cell Acute Lymphoblastic Leukemia. Int J Mol Sci. 2022;23(9).

23. Schmidt B, Brown LM, Ryland GL, Lonsdale A, Kosasih HJ, Ludlow LE, et al. ALLSorts: an RNA-Seq subtype classifier for B-cell acute lymphoblastic leukemia. Blood Adv. 2022 Jul 15;6(14):4093–7.

24. Beder T, Hansen BT, Hartmann AM, Zimmermann J, Amelunxen E, Wolgast N, et al. The Gene Expression Classifier ALLCatchR Identifies B-cell Precursor ALL Subtypes and Underlying Developmental Trajectories Across Age. HemaSphere [Internet]. 2023;7(9). Available from:https://journals.lww.com/hemasphere/fulltext/2023/09000/the_gene_expression_classifier_allcatchr.7.aspx

25. Hu Z, Jia Z, Liu J, Mao A, Han H, Gu Z. MD-ALL: an integrative platform for molecular diagnosis of B-acute lymphoblastic leukemia. Haematologica. 2024 Jun 1;109(6):1741– 54.

26. Li L, Xiao H, Wu X, Tang Z, Khoury JD, Wang J, et al. RanBALL: An Ensemble Random Projection Model for Identifying Subtypes of B-cell Acute Lymphoblastic Leukemia. bioRxiv. 2024 Jan 1;2024.09.24.614777.

27. Gu A, Schmidt B, Lonsdale A, Jalaldeen R, Kosasih HJ, Brown LM, et al. TALLSorts: a T-cell acute lymphoblastic leukemia subtype classifier using RNA-seq expression data. Blood Adv. 2023 Dec 13;7(24):7402–6.

28. Aplenc R, Meshinchi S, Sung L, Alonzo T, Choi J, Fisher B, et al. Bortezomib with standard chemotherapy for children with acute myeloid leukemia does not improve treatment outcomes: a report from the Children’s Oncology Group. Haematologica. 2020;105(7):1879.

29. Bolouri H, Farrar JE, Triche T, Ries RE, Lim EL, Alonzo TA, et al. The molecular landscape of pediatric acute myeloid leukemia reveals recurrent structural alterations and age-specific mutational interactions. Nat Med. 2018 Jan 1;24(1):103–12.

30. Fornerod M, Ma J, Noort S, Liu Y, Walsh MP, Shi L, et al. Integrative Genomic Analysis of Pediatric Myeloid-Related Acute Leukemias Identifies Novel Subtypes and Prognostic Indicators. Blood Cancer Discov. 2021 Nov 1;2(6):586–99.

31. Faber ZJ, Chen X, Gedman AL, Boggs K, Cheng J, Ma J, et al. The genomic landscape of core-binding factor acute myeloid leukemias. Nat Genet. 2016 Dec 1;48(12):1551–6.

32. de Rooij JDE, Branstetter C, Ma J, Li Y, Walsh MP, Cheng J, et al. Pediatric non–Down syndrome acute megakaryoblastic leukemia is characterized by distinct genomic subsets with varying outcomes. Nat Genet. 2017 Mar 1;49(3):451–6.

33. Umeda M, Ma J, Huang BJ, Hagiwara K, Westover T, Abdelhamed S, et al. Integrated Genomic Analysis Identifies UBTF Tandem Duplications as a Recurrent Lesion in Pediatric Acute Myeloid Leukemia. Blood Cancer Discov. 2022 May 5;3(3):194–207.

34. McNeer NA, Philip J, Geiger H, Ries RE, Lavallée VP, Walsh M, et al. Genetic mechanisms of primary chemotherapy resistance in pediatric acute myeloid leukemia. Leukemia. 2019 Aug 1;33(8):1934–43.

35. Newman S, Nakitandwe J, Kesserwan CA, Azzato EM, Wheeler DA, Rusch M, et al. Genomes for Kids: The Scope of Pathogenic Mutations in Pediatric Cancer Revealed by Comprehensive DNA and RNA Sequencing. Cancer Discov. 2021 Dec 1;11(12):3008–27.

36. Rusch M, Nakitandwe J, Shurtleff S, Newman S, Zhang Z, Edmonson MN, et al. Clinical cancer genomic profiling by three-platform sequencing of whole genome, whole exome and transcriptome. Nat Commun. 2018 Sep 27;9(1):3962.

37. Vaswani A, Shazeer N, Parmar N, Uszkoreit J, Jones L, Gomez AN, et al. Attention is all you need. Adv Neural Inf Process Syst. 2017;30.

38. Agarap A. Deep learning using rectified linear units (relu). ArXiv Prepr ArXiv180308375. 2018;

39. Zhao Y, Li MC, Konaté MM, Chen L, Das B, Karlovich C, et al. TPM, FPKM, or Normalized Counts? A Comparative Study of Quantification Measures for the Analysis of RNA-seq Data from the NCI Patient-Derived Models Repository. J Transl Med. 2021 Jun 22;19(1):269.

40. Robinson MD, Oshlack A. A scaling normalization method for differential expression analysis of RNA-seq data. Genome Biol. 2010 Mar 2;11(3):R25.

41. Law CW, Chen Y, Shi W, Smyth GK. voom: precision weights unlock linear model analysis tools for RNA-seq read counts. Genome Biol. 2014 Feb 3;15(2):R29.

42. Kolde R. Pheatmap: pretty heatmaps. R Package Version. 2019;1(2):726.

43. Panahi N, Hylkema T, Hawkins G, Manselle MK, Peplinski JH, Wallace LK, et al. LAMP5 Is an AML-Restricted Immunotherapeutic Target Enriched in High-Risk Acute Myeloid Leukemia. Blood. 2023 Nov 2;142(Supplement 1):1585–1585.

44. Talarmain L, Clarke MA, Shorthouse D, Cabrera-Cosme L, Kent DG, Fisher J, et al. HOXA9 has the hallmarks of a biological switch with implications in blood cancers. Nat Commun. 2022 Oct 3;13(1):5829.

45. Iacovino M, Chong D, Szatmari I, Hartweck L, Rux D, Caprioli A, et al. HoxA3 is an apical regulator of haemogenic endothelium. Nat Cell Biol. 2011 Jan 1;13(1):72–8.

46. Xiang P, Yang X, Escano L, Dhillon I, Schneider E, Clemans-Gibbon J, et al. Elucidating the importance and regulation of key enhancers for human MEIS1 expression. Leukemia. 2022 Aug 1;36(8):1980–9.

47. Scicchitano S, Giordano M, Lucchino V, Montalcini Y, Chiarella E, Aloisio A, et al. The stem cell-associated transcription co-factor, ZNF521, interacts with GLI1 and GLI2 and enhances the activity of the Sonic hedgehog pathway. Cell Death Dis. 2019 Sep 26;10(10):715.

48. Shafik R, Hassan N, El-Meligui Y, Shafik H. The Meningioma 1 (MN1) Gene is an Independent Poor Prognostic Factor in Adult Egyptian Acute Myeloid Leukemia Patients. Asian Pac J Cancer Prev. 2017;18(3):609–13.

49. McIntyre LM, Lopiano KK, Morse AM, Amin V, Oberg AL, Young LJ, et al. RNA-seq: technical variability and sampling. BMC Genomics. 2011 Jun 6;12(1):293.

50. Leek JT, Johnson WE, Parker HS, Jaffe AE, Storey JD. The sva package for removing batch effects and other unwanted variation in high-throughput experiments. Bioinformatics. 2012 Mar 15;28(6):882–3.

51. Goodfellow I, Pouget-Abadie J, Mirza M, Xu B, Warde-Farley D, Ozair S, et al. Generative adversarial networks. Commun ACM. 2020;63(11):139–44.

52. Chawla NV, Bowyer KW, Hall LO, Kegelmeyer WP. SMOTE: synthetic minority over-sampling technique. J Artif Intell Res. 2002;16:321–57.

53. Mumuni A, Mumuni F. Data augmentation: A comprehensive survey of modern approaches. Array. 2022 Dec 1;16:100258.

54. Anilkumar KK, Manoj VJ, Sagi TM. A review on computer aided detection and classification of leukemia. Multimed Tools Appl. 2024 Feb 1;83(6):17961–81.

